# Identification of a signature of evolutionarily conserved stress-induced mutagenesis in cancer

**DOI:** 10.1101/2021.04.17.440291

**Authors:** Luis H. Cisneros, Charles Vaske, Kimberly J. Bussey

## Abstract

The clustering of mutations observed in cancer cells is reminiscent of the stress-induced mutagenesis (SIM) response in bacteria. SIM employs error-prone polymerases resulting in mutations concentrated around DNA double-strand breaks with an abundance that decays with genomic distance. We performed a quantitative study on single nucleotide variant calls for whole-genome sequencing data from 1,950 tumors and non-inherited mutations from 129 normal samples. We introduce statistical methods to identify mutational clusters and quantify their distribution patterns. Our results show that mutations in both normal and cancer samples are indeed clustered and have shapes indicative of SIM. We also found that clusters of mutations colocalize more prevalently in normal samples than in cancer, suggesting loss of regulation leading to increased randomness in the mutational process during carcinogenesis.

## Introduction

Genomic instability is a well-known hallmark of cancer which manifests as higher than normal rates of genomic mutations. However, these mutations do not typically arise at uniformly random locations across the genome. Rather, they typically follow a non-uniform distribution resulting in mutational clustering ***Drake (2007***); ***Wang et al. (2007***); ***Chen et al. (2009***); ***Ye et al. (2010); Roberts et al. (2012); Nik-Zainal et al. (2012); Alexandrov et al. (2013); Kamburov et al. (2015); Nik-Zainal et al. (2016***). In its extreme form, this phenomenon is observed as *kataegis*, a process consisting of six or more mutations with inter-mutational distances of one kilobase (kb) or less ***Alexandrov et al. (2013); Nik-Zainal et al. (2016***).

Given that most genomic variants are either neutral or deleterious, the likelihood that randomly distributed mutations would result in gains in fitness is considered to be very low ***Ram and Hadany (2014***). However, concerted patches of mutations, particularly when occurring within specific genes, could lead to neo-functionalization and increased cellular fitness ***Drake (2007); Ram and Hadany (2014); Cortés-Ciriano et al. (2020***).

Previous work has shown that even though cancer samples typically exhibit many mutations outside of genes, clustered mutations are more enriched in genes relative to the intergenic spaces ***Cisneros et al. (2017); Supek and Lehner (2017***). In particular, mutation clustering in non-coding regions have been associated with structural changes that possibly cause elevated mutation rates but by themselves very rarely constitute oncogenic drivers ***Nik-Zainal et al. (2016); Rheinbay et al. (2020***).

Large mutational loads in human cancer has been associated with replication repair deficiency ***Campbell et al. (2017); Ma et al. (2018); Campbell et al. (2021***), and thus underlying defects in the DNA repair machinery are thought to lead to biases in the types and locations of passenger mutations and structural events acquired during the progression of cancer. These general ideas justify targeting DNA repair and checkpoint inhibitors in cancer therapies ***Murai (2017); Forment and O’Connor (2018); Ubhi and Brown (2019); Zhu et al. (2020***). Other studies have identified the action of the AID/APOBEC family of cytosine deaminases as well as the action of Pol-*η* as contributing mechanisms to the phenomenon of mutational clustering in cancer ***Lada et al. (2012); Roberts et al. (2013); Taylor et al. (2013); Supek and Lehner (2017); Buisson et al. (2019); Roper et al. (2019); Shi et al. (2020***). However, these processes explain only a subset of the mutational clusters observed.

In bacterial, stress-induced mutagenesis (SIM) occurs when DNA double strand break damage (DSB) happens in the context of sufficient cellular stress to initiate the SOS response ***McKenzie et al. (2000); Foster (2007); Janion (2008); Shee et al. (2012); Rosenberg et al. (2012***). SIM has been shown to increase mutation rates locally around DNA lesions as cells strive to adapt to a challenging environment ***Foster (2007); Rosenberg et al. (2012); Fitzgerald et al. (2017***). During DSB-mediated mutagenesis in bacteria, DNA repair switches from high-fidelity homologous recombination to a repair mechanism that relies on the error-prone DNA polymerase Pol IV, encoded by the gene *dinB*. The result of this mechanism is a spectrum of both single nucleotide variants (SNV) and copy number amplifications. The molecular signature of this process is clustering of SNVs around the DSB site, with an event frequency that decays with the distance and remains above background for up to one megabase ***Rosenberg et al. (2012); Shee et al. (2012); Fitzgerald et al. (2017***).

Considering the evidence of mutational clustering, as well as the evidence of intra-tumor chromosomal structural heterogeneity that characterizes many cancers ***Roschke et al. (2002***, 2003, 2005) prompted us to inquire whether a process comparable to bacterial SIM takes place during carcinogenesis. This idea was previously suggested by Fitzgerald, Xia, Rosenberg and colleagues ***Fitzgerald et al. (2017); Xia et al. (2019***). In fact, expression of adaptive mutagenesis has been shown in the context of the emergence of drug resistance, with evidence of down-regulation of mismatch repair (MMR) and homologous recombination (HR), and up-regulation of error-prone polymerases in drug-tolerant colorectal tumor cells ***Russo et al. (2019***). Furthermore, mTOR stress signaling has been shown to facilitate SIM in multiple human cancer cell lines exposed to non-genotoxic drug selection ***Cipponi et al. (2020***).

In this study, we investigate SNV distributions observed by whole genome sequencing of non-inherited mutations in normal samples and in a wide variety of solid tumors. We examine whether these SNV distributions provide evidence of mutational clustering, whether the molecular signal of SIM can be identified in the SNV distributions, and how they relate to clinical outcomes.

## Results

### Variant distribution is not uniform

We analyzed the patterns of mutational density across the genome for non-inherited mutations from 129 normal individuals in the Complete Genomics Indices database of the 1000 Genome Project (CGI data), as well as somatic mutations in 1,950 tumors from 14 different tissues in the Pan-Cancer Analysis of Whole Genomes database (PCAWG data; see Methods/Data 1 for details), and compared them with simulated patterns of *N*_SNV_ = 1000, 2500, 5000, 10000, 25000, and 50000 total uniformly distributed mutations (500 replicates each) following our theoretical null model (Poisson point process model, see Methods/Model 1).

First, for values of *x* binned in 15 kb-bins up to 150 kb, we measured the number of inter-SNV segments of length *x* observed in each sample as a function of its mutational load (Fig.1). Perhaps the most distinct feature of this relationship is that it is not monotonic. For all inter-SNV distance range cases, as the total number of mutations increased, the theoretical prediction of the number of segments proportionally increased as well, then peaked, followed by rapid decrease. This drop is due to a saturation effect: in a three billion-base genome, if the number of uniformly distributed SNVs exceeds 100,000 the expected inter-SNV distance falls under 30kb, and thus long inter-SNV intervals become unlikely (Fig.1(A)).

**Figure 1.**
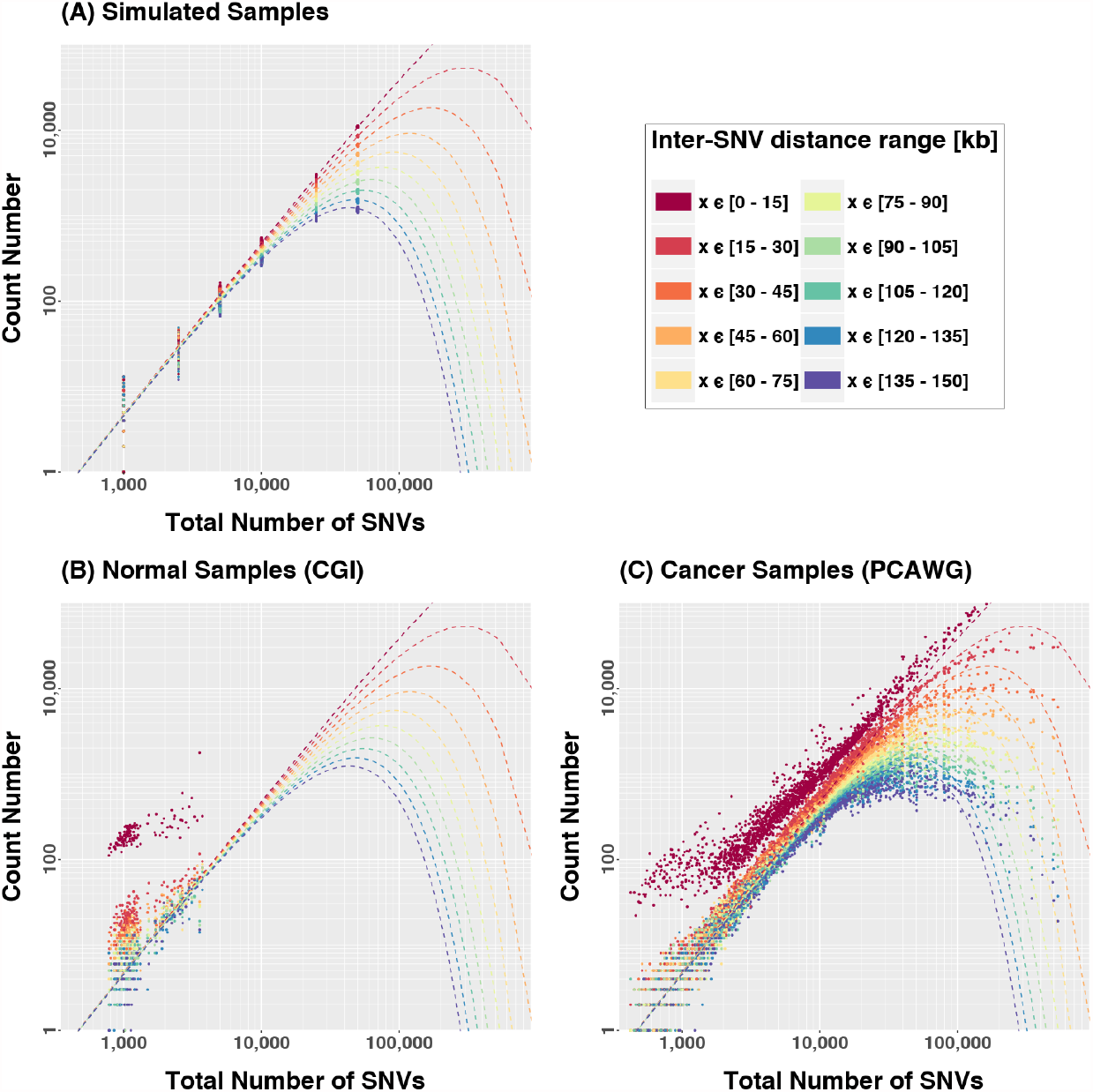
Count numbers of inter-SNV segments of different lengths as a function of the mutational load in (A) simulated, (B) normal and (C) cancer samples. Dashed lines are the theoretical predictions for a Poisson point process. Both normal and cancer cases show significant enrichment of small segments, indicating that mutations are typically closer than expected.

In both normal samples (Fig.1(B)) and cancer samples (Fig.1(C)), short *x*-segments were more frequently observed than expected from the null model, thus revealing a tendency of mutations to cluster together in genomic space. The effect was considerably stronger for lower mutational loads, particularly with *N*_SNV_ < 3000 where the number of short segments (*x* ≤ 15 kb) can be over an order of magnitude larger than expected. On the other hand, segments become progressively less over-represented as they get longer: *x* ∼ 75 kb segments appeared to have the expected frequency, and segments with *x* > 135 kb are typically under-represented.

Interestingly, in cancer samples the over-representation of small segments was prevalent even for large mutational loads. This effect was compensated for by an under-representation of moderate to long intervals, yet for very large mutational loads (*N*_SNV_ > 100, 000) long intervals were more frequent than expected. This suggests that some regions of the genome might be protected from acquiring mutations, which is manifested in the form of unexpectedly long conserved (mutation free) segments. These features were consistent across all samples and are evidently not associated with number fluctuations or sampling bias, since the dispersion in 500 simulated replicates cannot account for them (Fig.1(A)). From this analysis we conclude that mutations in both normal and cancer samples tend to form clustered groups.

We then looked at the number of groups, with “group” defined as a set of contiguous SNVs with inter-SNV distances *x* ≤ *D*^⋆^ (*D*^⋆^ = 15 kb) as a function of the mutational load. We deemed these groups *tuples*, while a *singleton* is a mutation that is not grouped (i.e., a 1-tuple).

The numbers of tuples and singletons for simulated, normal and cancer data are shown in Fig. 2(A). The most salient feature is that for low mutational loads the number of singletons was significantly under-represented with respect to the theoretical expectation, while tuples were typically over-represented (see Appendix section 1 for detailed distributions for different tuple sizes). The number of tuples was particularly high for samples with *N*_SNV_ < 3000, at which a uniformly random process would very rarely lead to any proximal mutations with *x* ≤ *D*^⋆^. Specifically, at *N*_SNV_ ∼ 1000 only a handful of tuples are expected, yet dozens to hundreds were typically observed in both cancer and normal samples. As the total number of cancer mutations increased, the distributions approached the predicted curve, but then departed again (Fig. 2(A)). For large mutational loads the relationship between tuples and singletons with respect to the expected behavior is indeed inverted, supporting the idea that certain regions in the genome are protected from accumulation of clustered mutations. Those regions would commonly carry singletons and rarely contain tuples as would be expected in a uniform random process.

**Figure 2.**
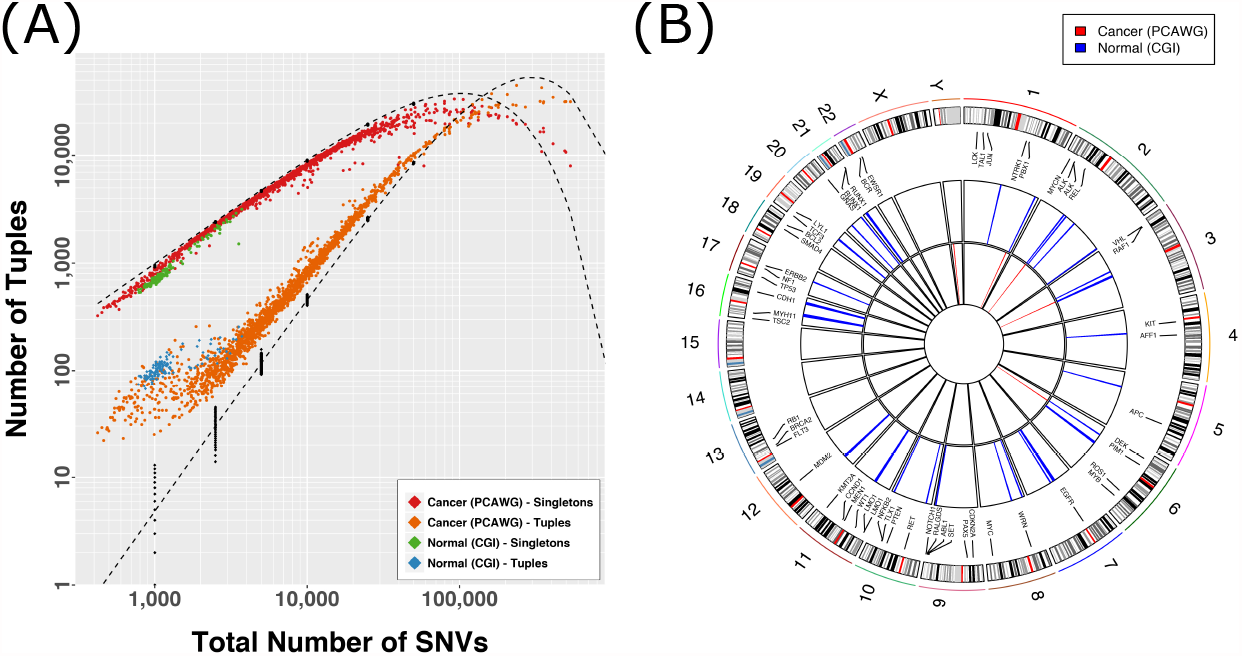
(A) Observed number of tuples and singletons as a function of the total mutational load. A tuple is a set of consecutive mutations with inter-event distance *x* ≤ 15 kb. A singleton is a mutation farther than 15 kb from any other mutation (1-tuple). Black dots are simulated data, dashed lines are the expected curves according to Poisson statistics. (B) Susceptible regions for samples with *N*_SNV_ < 5100, defined as genomic regions that overlap with tuples in at least 8.8% of the normal samples (blue) and 3.5% of the cancer samples (red) (percentages based on the square root of the number of samples in each set). These regions are evidently more common in normal than in cancer samples.

Altogether, these observations demonstrate that the mutational rate is a heterogeneous property of the genome, and thus likely a regulated and constrained process. We hypothesize two non-mutually exclusive ways to generate such mutational patterns:

1. The mutational process is modulated. This means that it can be modeled as a non-uniform Poisson process with location-dependent rate *λ*(*g*), with *g* the genomic location. This modulation can be an effect of differential DNA repair efficiencies along the genome, conservation of certain genomic regions, topological domains, folding/packing molecular properties, or other sequence-dependent processes.
2. Mutations are inter-dependent events, entailing either nucleation of mutations during subsequent DNA replications (i.e., mutations induce new, proximal errors during DNA translation) or a process in which events happen together as a single burst of proximal mutations.

If the mutational process is dependent on genomic location, then tuples locations would be prevalent across samples. We compared the tuple locations across samples for which tuple enrichment was most obvious (Fig. 2(A)): *N*_*s*_=129 normal samples and *N*_*s*_=784 cancer samples with *N*_SNV_ < 5100. To this end, we identified all regions in the genome containing tuples in at least 8.8% of the normal samples and 3.5% of the cancer samples (these fractions are picked to provide confidence that the observation is above the Poisson-counting error statistic). For normal samples (Fig. 2(B)) we found 128 overlapping regions, some containing tuples in as many as 30% of the samples. Many of these regions are located close together, as shown in Fig. 2(B), none of them were longer than 30 kb, and about a quarter of them involved single base mutations. In contrast, cancer samples had few overlaps. We observed only 19 regions grouped into five distinct ranges (see Fig. 2(B)): a ∼ 117-kb region in chromosome 6 associated with the human leukocyte antigen (HLA) complex, which contained tuples in 7% of the samples; two ∼ 1-kb regions in chromosomes 2 and 3 in 3.5% of the samples; a single point mutation in chromosome 1, associated with the zinc finger protein ZNF678, overlapping in ∼ 4% of the samples; and a half-kb region in chromosome Y with 11% overlap in the 479 male samples.

These results suggest that both processes in our hypotheses could be at play. Given that cancer samples display such poor overlap of the clustering pattern, the close proximity of mutations is possibly driven by a process in which events are not necessarily independent from each other, perhaps occurring simultaneously, yet otherwise distributed mostly randomly across the genome (and therefore not consistent between samples). In contrast, the stronger overlap between normal samples suggests that non-inherited mutations in normal tissue are at least partially driven by a location-specific and/or sequence-specific process, quite possibly sculpted by evolution and regulated across the genome.

### Quantification of Cluster Shapes

Previous work demonstrates that SNVs cluster together in both normal tissues and cancer, and the sequence contexts of both the reference and mutant calls can be used to infer mechanism ***Roberts et al. (2012***). The association of APOBEC cytosine deaminases with clusters is well established ***Lada et al. (2012); Burns et al. (2013); Taylor et al. (2013); Roberts et al. (2013***), but it only accounts for at most 50% of the clusters observed ***Roberts et al. (2013***). Furthermore, nothing in the mechanism of APOBEC suggests a characteristic shape of the mutational clusters. In contrast, the SIM response of bacteria, mediated by the Pol IV polymerase (encoded by *dinB*), leads to a clustering pattern in which more SNVs are found at the center of the cluster than at the edges ***Shee et al. (2012***). We define clusters as statistically unlikely tuples of size *n* ≥ 3, according to a negative binomial test (see Methods 1 for details). We then measured the number of clusters per sample, and their shapes, as a function of total mutational load among non-inherited mutations and somatic mutations in cancer samples.

Simulated data showed that as the total number of uniform random mutations increases, we expect to see the number of clusters and the fraction of SNVs that are in those clusters increase as well (Fig. 3(A)-(B)). We note that, in agreement with observations presented above, no clusters were observed in simulated data with mutational loads *N*_SNV_ < 2500, and a mean of only four clusters per genome was detected in samples with 2500 mutations. This indicates that in a uniform random process, at least several thousand mutations are required to expect any measurable clustering. In contrast, both non-inherited mutations in normal tissue and somatic mutations in cancer show extensive clustering when the mutational burden is that low (Fig 3(A)). On the other hand, for very large numbers of mutations we observed the expected saturation in the number of clusters.

**Figure 3.**
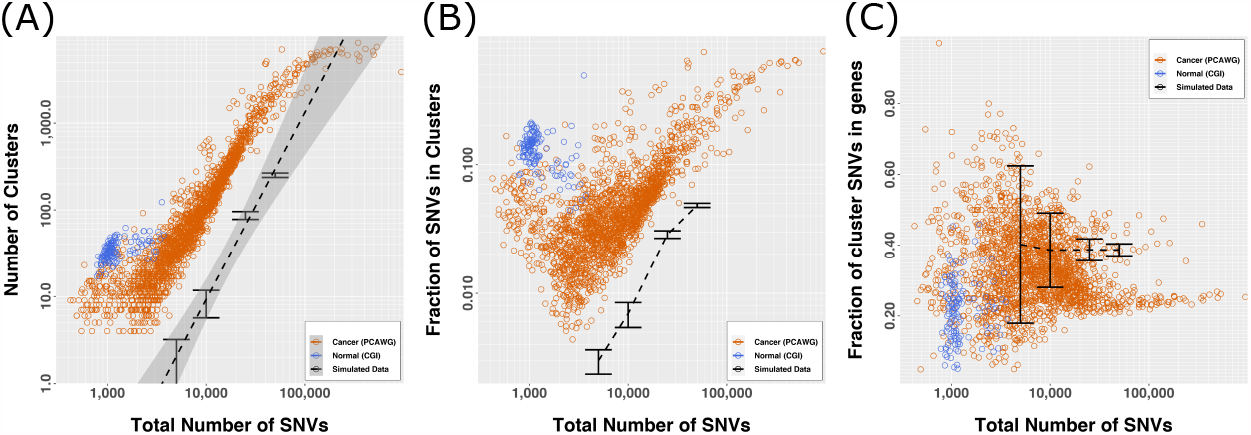
(A) Number of clusters per sample as a function of mutational load. The dashed line is the best fit for simulated data. Clustering is clearly more frequent than expected. The plateau at larger mutational loads is related to the limit at which the average inter-mutation distance approaches 30 kb, which produces many statistically likely tuples and thus fewer clusters. (B) Fraction of SNVs in clusters versus *N*_SNV_. The fraction of SNV in clusters increases with the number of mutations, suggesting that as more mutations are accumulated in the genome they are preferentially placed in clusters. (C) Fraction of cluster SNVs in genes versus *N*_SNV_, showing that clusters do not typically overlap with genes for high *N*_SNV_.

Figure 3(B) shows that as more mutations accumulate in cancer samples, a larger fraction of those mutations is preferentially placed in clusters. This raises the question of whether the clustering process itself is somehow implicated in the mechanism driving cancer mutations, in a sort of positive feedback loop or nucleation process. Another interesting observation is that the fraction of SNVs in clusters is about five times higher in normal samples than in to cancer samples with the same mutational load - a load for which we don’t expect any clustering at all under our null hypothesis. This indicates that the mutational process in normal samples is in fact driven by a mechanism that favors close proximity of variations and is likely restricted to susceptible genomic regions (Fig.2(B)).

In looking at whether the SNVs were located in genes or in intergenic regions, the null hypothesis predicts that as the number of SNVs increases, the proportion of SNVs located in genes converges to about 40% and then remains constant (Fig.3(C)). In normal samples, private non-inherited mutations oscillate around 37% (range 31.3%-42.8%) of SNVs localizing within genes, while cancer was defined by a large amount of variability that converged to about 25% of SNVs located in genes.

In agreement with our previous study ***Cisneros et al. (2017***), SNVs in clusters in cancer samples are preferentially excluded from genes (Fisher’s exact, Odds Ratio (OR) = 0.6002, 95% CI =0.5992-0.6013, p-value < 2.2×10^−16^). When we looked specifically at the position of clusters within genes by counting SNVs that overlap genes versus those that do not, we observed a slight enrichment for SNVs overlapping the 3’-end of genes that are in clusters compared to SNVs that are not in clusters (Fisher’s exact, OR = 1.024, 95% CI = 1.021-1.027, p-value =3.067×10^−57^), confirming previous observations by Supek and Lehner ***Supek and Lehner (2015***, 2017).

To evaluate the shapes of the clusters, we introduced the Stress-Introduced Heterogeneity (SItH) score (see Methods section and Fig. 6). The SItH score was computed both on individual clusters (cluster SItH) and over all clusters in a tumor (overall SItH). In simulated data, increasing the number of SNVs led to decreasing overall SItH scores (Fig. 4(A)) and produced a negative sigmoid shape for the variability in cluster SItH measured by the inner-quartile range (IQR) (Fig. 4(B)). The overall SItH score was higher in cancer samples than in normal samples. In comparison to the simulated data, overall SItH scores of both cancer and normal samples were higher at the extremes of mutational burden and lower in the mid-range of total SNV count (Fig. 4(A)). Moreover, cancer samples showed a greater diversity of SItH scores than predicted under the assumption of random uniform mutations or compared to normal (Fig. 4(B)). Interestingly, the diversity of cluster shapes reaches a plateau at mutational loads corresponding to higher than expected overall SItH scores. Combined, these results suggest that the mutational clustering in cancer is complex and likely driven by multiple mechanisms simultaneously.

**Figure 4.**
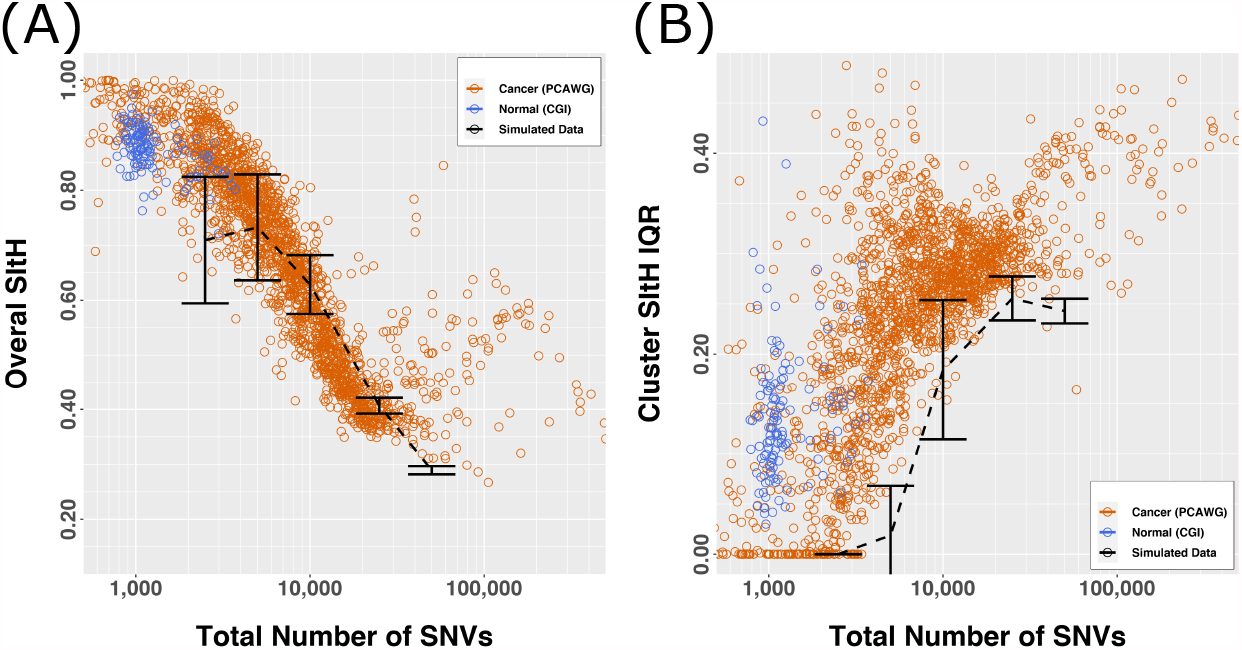
SItH scores by number of SNVs. (A) Overall SItH score as a function of mutational load. (B) Inner-quartile range (IQR) of cluster SItH scores as a function of mutational load.

**Figure 5.**
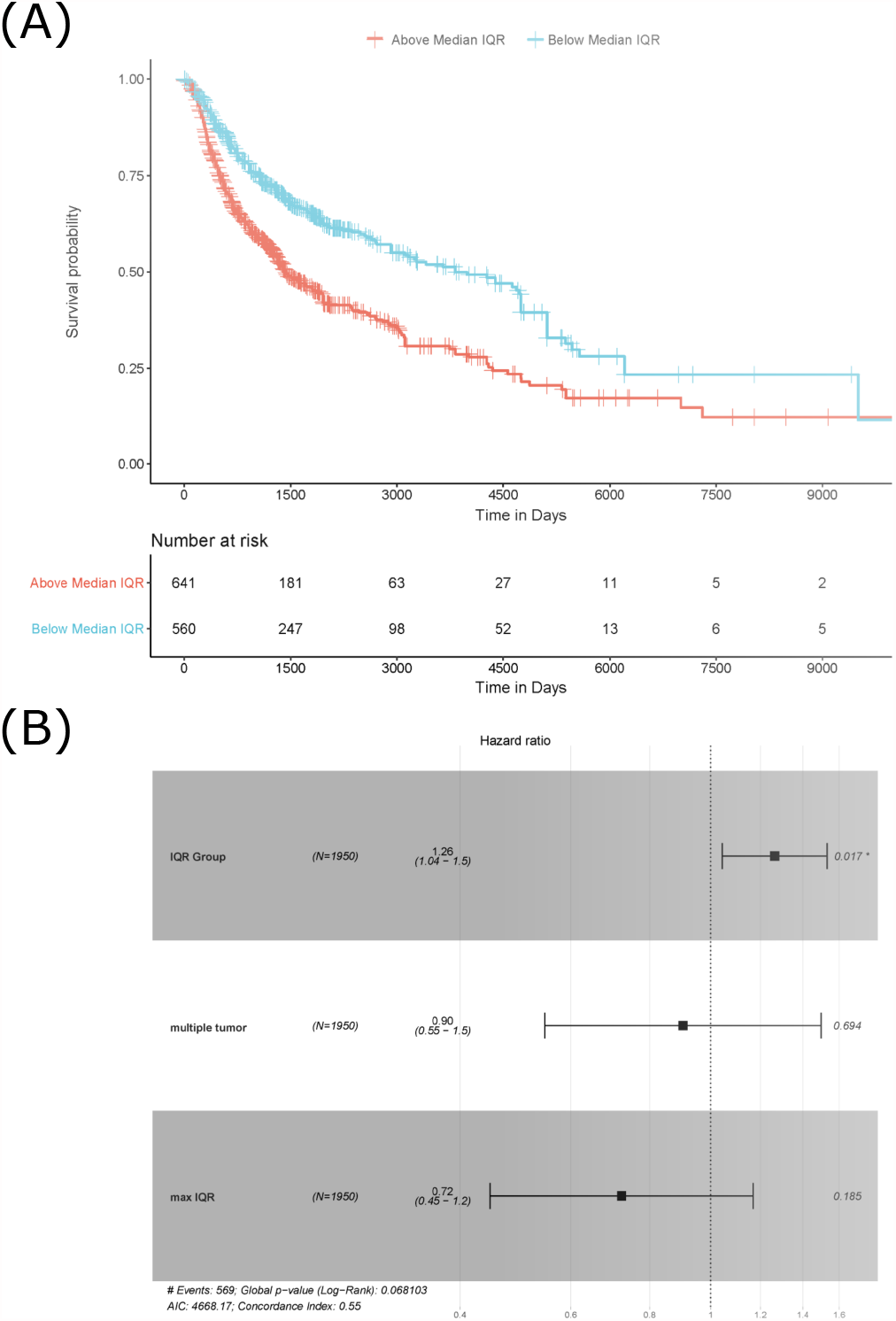
Survival difference based on SItH IQR being above or below the median score. a) Kaplan-Meir curves for tumors with cluster-level SItH IQR above and below the median SItH IQR for 1895 tumors. b) Results from the Cox proportional hazard analysis. Survival data from 1,950 tumors from 14 different cancer types. Hazard ratio for IQR group was controlled for maximum IQR value, tissue of origin, and multiple tumor samples for the same donor.

**Figure 6.**
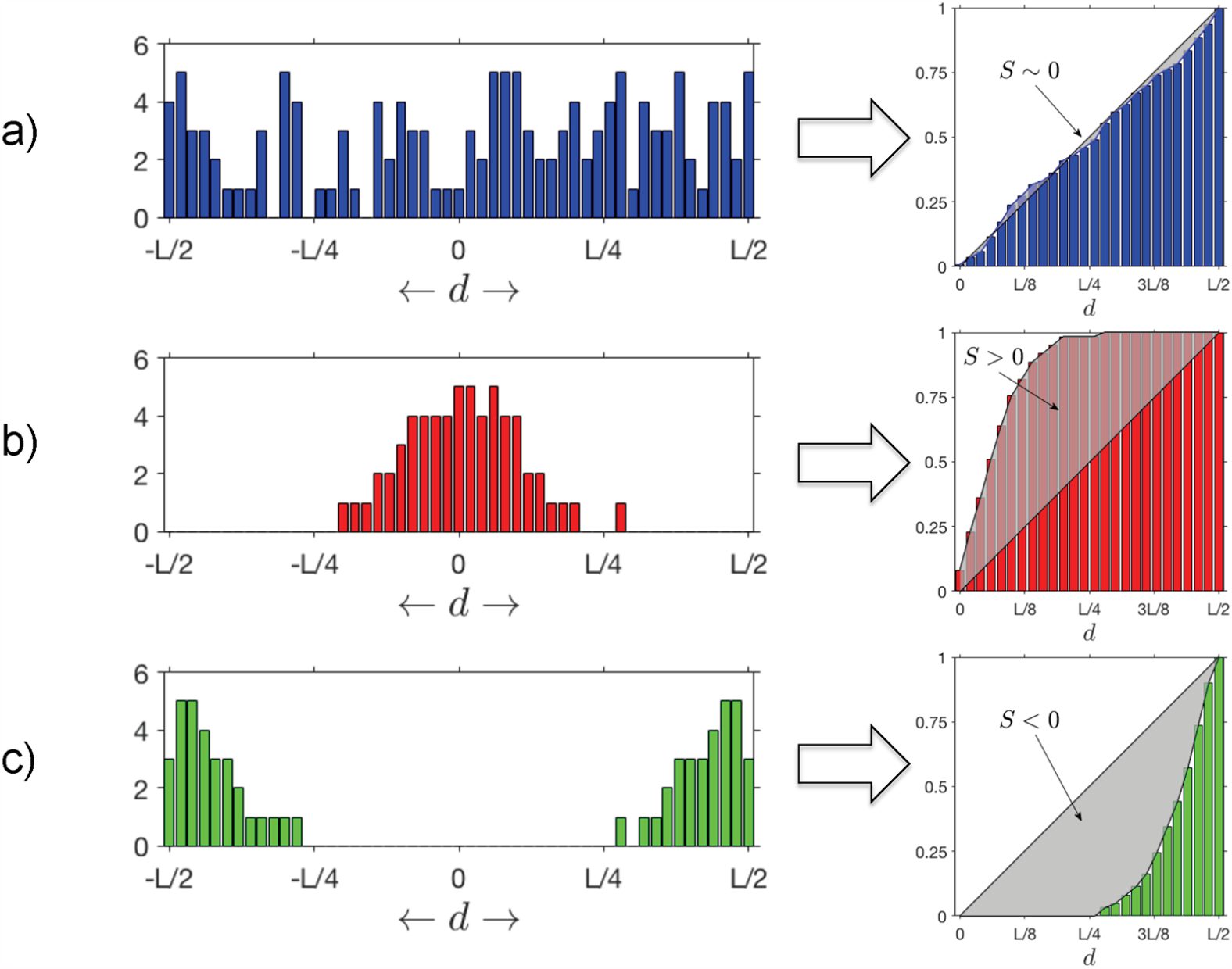
SItH Scores. Different outcomes for distributions of mutations as a function of the distance d to cluster centroids. (A) When *S* ∼ 0, uniformly distributed mutations yield a linear cumulative distribution. (B) *S* > 0 signifies a bell-shaped distribution of mutations around the centroid. (C) *S* < 0 signifies a distribution of mutations that increases with the distance to the centroid.

### Survival Analysis

A key characteristic of SNV clusters that result from SIM mechanisms is a decay in the frequency of incidental SNVs as a function of distance from the DSB that triggered error-prone repair response ***Shee et al. (2012***). We postulated that a more positive overall SItH score reflects a greater contribution of SIM to the mutational landscape of the tumor. Therefore, SItH provides a measure of the evolutionary response, or the adaptive capacity, of a tumor to a source of stress, such as chemotherapy. Overall SItH scores ranged from 0.145 to 0.999 (Fig. 4(A)) and varied significantly by organ site and whether the tumor was one of multiple tumors from a single donor (ANOVA, organ site, F = 136.70, p< 2.2×10^−16^; multiple tumor, F = 3.07, p = 0.0799; maximum SItH Score, F = 16.14, p = 6.098×10^−5^).

To determine the relationship between SItH scores and clinical outcome, we conducted Cox proportional hazard analysis of the overall SItH score as well as the IQR of the cluster SItH. The model for overall SItH is specified as follows:

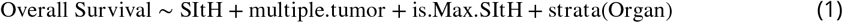

where the data analyzed were either primary tumors or the group of metastases and recurrences. For IQR of the cluster SItH the model is:

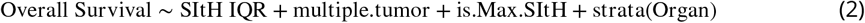

After controlling for organ site and multiple tumor status, we found that overall SItH scores predict patient survival, with different effects depending on whether the sample was a primary tumor or from a metastasis or recurrence. For primary tumors, more positive overall SItH scores predicted better patient survival (Cox Proportional Hazard Regression (CPHR), Hazard Ratio (HR) = 0.4516, 95% CI: 0.2274 -0.8968, p=0.0231). However, when the recurrences and metastatic tumors were considered as a group, more positive overall SItH scores predicted worse survival, with a HR of 14.84 (CPHR, 95% CI: 1.934-113.876, p= 0.00947). In looking at the diversity of SItH scores on a cluster basis, the type of tumor sample was no longer relevant. Wider IQR of cluster-level SItH scores was associated with worse survival, with a HR of 5.744 (CPHR, 95% CI: 1.824 -18.09, p= 0.00283). We then examined whether there was a difference in survival between patients with SItH IQRs above or below the median SItH IQR, as clinical translation will likely require creating a cut-off value above which one would predict poor prognosis. As seen in Fig. 5, there is a significant difference in survival, even after accounting for the baseline differences in survival by tissue of origin (CPHR, HR = 1.26, 95% CI: 1.043-1.531, p= 0.0168).

## Discussion

Our study provides evidence that a signature of stress-induced mutagenesis, characterized by clustering of SNVs, is widespread across multiple cancer types. Both the strength of SIM and the diversity of mutational processes within a tumor are expected to impact disease outcome ***Andor et al. (2016***). Our results show an association of both overall cluster shape (overall SItH) and increased cluster shape diversity (SItH IQR) with patient survival. We submit that SItH IQR predominantly represents the amount of time SIM has been active during carcinogenesis and clonal diversification, while overall SItH represents the ratio of the intensity of SIM relative to other mutational processes. Our work shows that an increase in mutational load leads to increases in both cluster size and the percentage of SNVs involved in clusters, but only up to a point. In tumors with high mutational burdens, the number of clusters, the genomic distance covered by clusters, and the number of SNVs contained within a cluster all level out. This implies that under high mutational burden the variations in mutation density across the genome flatten out, likely due to alterations in DNA repair pathways, such as a loss of mismatch repair ***Supek and Lehner (2017); Campbell et al. (2017***) that obscures the detection of clusters.

The influence of intra-tumor diversity on clinical outcome is an area of active investigation. Evidence from measures of clonal diversity and copy number diversity are associated with both worse outcome and therapeutic response ***Andor et al. (2016); Davoli et al. (2017); Roh et al. (2017); Dagogo-Jack and Shaw (2017); Turajlic et al. (2019); Ben-David and Amon (2019***). Cancer must balance the introduction of genomic rearrangements that contribute to cellular diversity with a sufficient level of genome stability to avoid a genomic error catastrophe. Our results are consistent with this idea, in that large positive overall SItH scores in primary tumor samples are associated with better patient survival. The SItH IQR represents a measure of mutational heterogeneity that ties intra-tumor diversity to an underlying evolutionarily conserved process in response to cellular stress. In other words the SItH IQR is a measure of the heterogeneity of adaptive strategies within a patient. This diversity manifests as a broad ensemble of mutational cluster shapes within a tumor, driven by the heterogeneity in mutational processes to generate genomic diversification. This in turn increases the substrates available for broad phenotypic plasticity, including transcriptional responses. Such responses have been shown to be important in the rapid acquisition of resistance to Doxorubicin ***Wu et al. (2015***). In this case high diversity results in a direct survival advantage for the tumor, allowing it to respond to a wider range of stresses and leading to poorer outcomes for patients.

Others have proposed mechanisms for clustered mutations in cancer ***Roberts et al. (2012); Lada et al. (2012); Burns et al. (2013); Taylor et al. (2013); Roberts et al. (2013); Supek and Lehner (2017***). In particular, Supek and Lehner showed that Pol-*η*, a TLS polymerase closely related to Pol IV, is involved in the generation of clustered mutations that preferentially locate at the 3’-end of active genes ***Supek and Lehner (2015***). Although our method uses a broader definition of mutational clusters than previously proposed (***Shee et al. (2012); Fitzgerald et al. (2017***)) we were able to confirm this key finding.

An open question that remains is whether the clusters we, and others, have detected arose from singular events reflective of bursts of mutational activity, or were accumulated over time. The latter scenario would identify distinct regions of the genome prone to mutation. Measuring allele fraction has been suggested as one way to address this question. However, the limited precision of most allele fraction measurements prevents the accurate discrimination of varying degrees of heterogeneity across a tumor. For example, the 95/95 binomial tolerance interval for a true allele fraction of 0.5 at a read depth of 60x ranges from 0.25 to 0.75 (see Appendix 2). This interval represents the bounds in which we are 95% confident that 95% of the measurements of a true allele fraction of 0.5 will lie. If we have a cluster where the allele fractions of multiple SNVs all fall within this range, we cannot rule out whether these actually represent a true allele fraction of 0.5 and therefore all come from the same event. Experimental evidence in mammalian systems leading to cluster formation is necessary to answer this question. This is an important study to pursue as the strategies one might propose for influencing mutational patterns with impact on clinical outcomes will depend on whether the target is the mutational process itself or the regions of the genome being acted upon by the mutational process.

## Conclusions

Cancer is notorious for outsmarting physicians. To make progress, we need to factor in how cancer cells evolve and adapt in the face of the challenges of medical treatment. A deeper understanding of the mechanisms of mutation and adaptation in cancer is therefore an essential pre-requisite for improving patient outcomes. Stress-induced mutagenesis, an ancient and evolutionarily conserved adaptive mutation mechanism well-characterized in *E. coli*, comprises some part of the genomic instability seen in cancer and contributes to the ability of the tumor to evolve resistance to therapy ***Fitzgerald et al. (2017***). We have described a way to quantify this biological response and shown that SIM has a strong association with poor prognosis. Further investigations into the process of SIM in cancer should lead to better patient outcomes by giving clinicians a metric by which they can tailor treatments to regulate tumor progression and minimize the risk of triggering an aggressive evolutionary response.

## Methods

### Null model - Uniform random mutations

A uniform distribution of point mutations can be modeled as a Bernoulli process. Because the total number of mutations is typically a lot smaller than the number of nucleobases in the genome the expected inter-event distance can be approximated as an exponential function. This distribution is equivalent to the expression for the waiting time distribution in a Poisson process ***Cinlar (1975***) or the survival density function in a constant hazard process ***Moore (2016***).

The probability of observing an inter-event distance *x* is given by the density function:

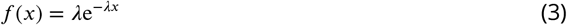

where *λ* = N_SNV_/*L* is the mutational rate, N_SNV_ is the total number of SNV mutations, and *L* is the length of the genome. For a given distance *D*, the probability of *x* ≤ *D* is the *cumulative waiting time function*:

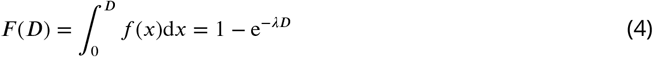

and the probability of *x* > *D* is the *survival function*:

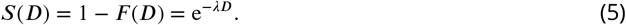

The probability associated with the range of interval lengths [*d*_*i*_, *d*_*i*_ + *D*_bin_] is

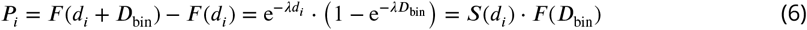

and the expected number of intervals with length in this range is

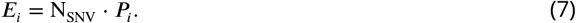

We define a *n*-tuple as a set of *n* consecutive mutations that are closer than *D*^⋆^ = 15 kb. By definition all tuples are separated by intervals *x* > *D*^⋆^ from each other. In particular, 1-tuples, or *singletons*, are SNVs that are farther than *D*^⋆^ bases from their closest neighbors.

From Eq. (6) the probability of *x* ≤ *D*^⋆^ is *F* ^⋆^ = 1 − e^−*λD*⋆^ and the probability of *x* > *D*^⋆^ is *S*^⋆^ = e^−*λD*⋆^. The expected total number of tuples can be estimated as:

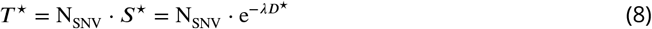

The total number of mutation events in tuples can be estimated as *E*^⋆^ ∼ N_SNV_ ·*F* ^⋆^. This identity is not exact because of edge effects, particularly location differences are calculated on each chromosome independently (not including Y), and therefore the total number of observed inter-mutation intervals is N_SNV_ − 25.

Following these definitions, the probability of observing a *n*-tuple can be written as the combination of probabilities:

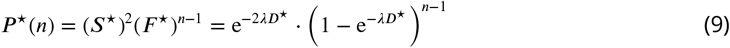

Thus the expected number of *n*-tuples is *N*^⋆^(*n*) = N_SNV_ · *P* ^⋆^(*n*) and the probability mass function of *n*-tuples is

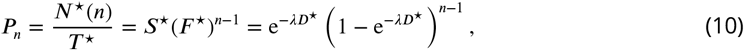

which is equivalent to the binomial mass function of the first order *P*_*r*_

### Data

We obtained variant calls for normal and cancer samples from public repositories. For non-inherited mutations in normal tissue, we used whole genome sequencing data (WGS) from the Complete Genomics Indices database in the 1000 Genomes Project ***The 1000 Genomes Project Consortium (2015***)(release 20130502, see Supplementary Materials Table 7 in Cisneros, et al. ***Cisneros et al. (2017***) for a list of donors). These data have an average genome coverage of 47*X*. The variant call tables (VCF) of 129 trios were analyzed using the vcf_contrast function from the VCFTools analysis toolbox to compare each child with the two corresponding parents. The resulting potential novel variants were then filtered such that the child and both parents must be flagged as PASS (i.e., the variant passed all filters in the calling algorithm); the child must have a read depth of at least 20; and the alternative (aka novel) allele frequency was ≥ 0.35. For cancer samples, we analyzed simple somatic mutations and corresponding clinical data from the PCAWG coordinated WGS calls for 1,950 tumor samples from 1,830 donors representing 14 different primary sites ***Campbell et al. (2017***). Somatic variants for all data sets were classified as previously published ***Cisneros et al. (2017***).

For our theoretical data, we generated 500 replicates for eight groups of simulated mutations defined by their total mutational load (*N*_SNV_ = 500; 1, 000; 2, 500; 5, 000; 10, 000; 25, 000; 50, 000; 100, 000). We modeled a uniform, random distribution of SNVs across the genome as a one-dimensional Bernoulli Process.

The number of events in a region of size *X* is a random variable with a probability mass function that can be approximated as a Poisson distribution:

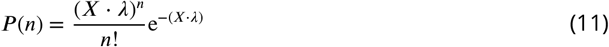

where *λ* = (*N*_SNV_/*L*) the total mutational rate and *L* is the genome length.

In order to characterize the clustering of genomic mutations we defined a tuple as a set of consecutive SNVs such that all its inter-event distances are shorter than *D*^⋆^ = 15 kb. According to Poisson statistics (per Eq.(10)) the expected number of n-tuples in a sample with *N*_SNV_ mutations is given as

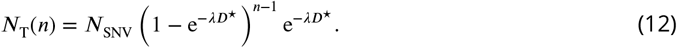

### Detection of mutation clusters

A group of SNVs is deemed a “cluster” if it is a tuple of at least 3 variations and the probability of finding it by random chance is less than 1% according to the negative binomial regression given by the total rate of observed mutations in the genome. In other words, the particular group of variations is statistically unlikely to happen in the background given by the mutational load of the sample.

In principle a given tuple might not satisfy the negative binomial test condition but part of it could (e.g. a tuple with higher concentration of mutations in one end). Our method accounts for this possibility so that only the statistically significant portion of a tuple is called a “cluster”.

For each WGS sample in our database, all possible clusters were identified and the “center of mass” (genomic location of cluster centroid) in each case was calculated, along with other properties such as start and end locations, length, and size (number of variations) ***Cisneros et al. (2017***).

### Detection of cluster shape

We treated cluster centroids as likely locations of the DSBs that induced the accumulation of variations. Therefore, the expected signature for stress-induced mutagenesis should be evident as a concentration of mutations around these centroids that decays with distance. Thus, for each cluster *i* we computed the cumulative distribution of SNV events *F*_*i*_(*X*) as a function of the distance *X* from the cluster centroid up to 250 kb and in both the 3′ and the 5′ directions. By aggregating all cumulative distributions observed in each sample we generated a representative overall curve *F* (*X*) = ∑ *F*_*i*_(*X*) that conveys the probability of finding a mutation at a given distance from the cluster center. If the distribution of SNV events is uniformly random (and therefore does not typically decay) then *F* (*X*) is expected to increase proportionally with *X*. This assumption gives us a background of mutations against to which we can compare the observed distribution pattern. It is important to note that this definition is itself independent of the definition of clusters. By construction, if the background distribution is uniform as assumed, then we should not observe clusters at all since they are statistically unlikely by random chance. In order to define a useful score, we normalize *X* by 250 kb and *F* by the number of events closer than 250 kb, thus mapping all cluster-associated cumulative distribution curves to a unit box:

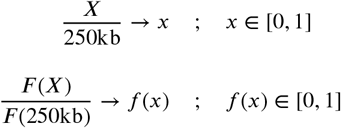

If the null hypothesis were correct for these events, *f* (*x*) = *x*. We define a measure of the degree of deviation from the null hypothesis by integrating the difference between the normalized cumulative distribution *f* (*x*) and the expected value *x* as follows:

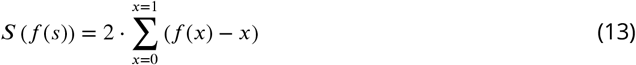

The value of *S* is a signed statistic with range *S* ∈ [−1, 1] (see Figure 6). As *S* approaches one, smaller windows close to the origin (i.e., cluster center) contain more events than expected from a random uniform distribution, indicating that SNV events concentrate near the center of the clusters and sharply decay with the distance. A negative *S* value indicates that the events are typically depleted from the center and concentrated on the edges of the cluster, and *S* values close to zero indicate that events are mostly uniformly distributed across the 250 kb interval length, supporting the null hypothesis. We call this the **Overall Stress Introduced Heterogeneity** (SItH) score of the distribution of somatic SNVs, and use it to represent the typical cluster geometry in a sample. Following the same definition for individual clusters we can estimate a **Cluster SItH** score using the function *F*_*i*_(*X*) instead of *F* (*X*), thus leading to *S*_*i*_ = *S*(*f*_*i*_(*x*)). This definition is statistically less robust than the overall measure but allows us to assess the diversity of behaviors in clusters within a sample. We do this by estimating the quartile statistics on the ensemble of *S*_*i*_ values for each sample.

### Effects of the maximum inter-SNV distance on the definition of clusters

The specific value *D*^⋆^ = 15 kb was chosen because it is an adequate balance between signal and noise. If *D*^⋆^ is too small, very few clusters are found unless the total number of mutations is very large. However, the restrictive condition given by a smaller *D*^⋆^ does yield more concentrated clusters with small dispersion as measured by the SItH IQR. On the other hand if *D*^⋆^ is too large many clusters are found and the noise level is higher. In this case there is more room for different configurations of clusters, producing larger IQR values. Low to moderate mutational loads typically have smaller SItH scores since clusters tend to be more uniform (i.e., less peaked), but samples with large mutational loads exhibit saturation effects that limit the number of clusters (i.e., if tuples are common then they are not clusters). There is therefore a trade-off between the effects on smaller and larger mutational loads. In order to find a good signal-noise balance, we ran our analysis with eight different values of *D*^⋆^. Tables 1 and 2 show the correlations between the overall SItH score and SItH IQR in cancer samples. Based on these results we conclude that *D*^⋆^ = 15 kb is a good choice; result values correlate well with cases in both ends, indicating that this parameter captures well the signal for both small and large mutational loads without too much compromise in quality.

**Table 1.** Correlation of overall SItH scores for different *D*^⋆^ values with PCAWG data.

| SlitH | 1kb | 2kb | 5kb | 10kb | 15kb | 20kb | 25kb | 50kb |
| --- | --- | --- | --- | --- | --- | --- | --- | --- |
| 1kb | 1 | 0.988498379 | 0.953254764 | 0.913062667 | 0.885251282 | 0.872055507 | 0.86602581 | 0.890076618 |
| 2kb | 0.988498379 | 1 | 0.979158317 | 0.944003381 | 0.914886845 | 0.896847418 | 0.884576285 | 0.885092543 |
| 5kb | 0.953254764 | 0.979158317 | 1 | 0.981798123 | 0.95732091 | 0.937928221 | 0.92095692 | 0.885284312 |
| 10kb | 0.913062667 | 0.944003381 | 0.981798123 | 1 | 0.988106173 | 0.973637963 | 0.957521878 | 0.900404232 |
| 15kb | 0.885251282 | 0.914886845 | 0.95732091 | 0.988106173 | 1 | 0.992613883 | 0.980773228 | 0.919694508 |
| 20kb | 0.872055507 | 0.896847418 | 0.937928221 | 0.973637963 | 0.992613883 | 1 | 0.993667498 | 0.939087871 |
| 25kb | 0.86602581 | 0.884576285 | 0.92095692 | 0.957521878 | 0.980773228 | 0.993667498 | 1 | 0.957075149 |
| 50kb | 0.890076618 | 0.885092543 | 0.885284312 | 0.900404232 | 0.919694508 | 0.939087871 | 0.957075149 | 1 |

**Table 2.** Correlation of SItH IQR scores for different *D*^⋆^ values with PCAWG data.

| SlitH IQR | 1kb | 2kb | 5kb | 10kb | 15kb | 20kb | 25kb | 50kb |
| --- | --- | --- | --- | --- | --- | --- | --- | --- |
| 1kb | 1 | 0.938197026 | 0.838362333 | 0.78391552 | 0.737391598 | 0.688942755 | 0.65370155 | 0.445513598 |
| 2kb | 0.938197026 | 1 | 0.906115983 | 0.830270283 | 0.774522805 | 0.721045756 | 0.680923471 | 0.454410934 |
| 5kb | 0.838362333 | 0.906115983 | 1 | 0.921681646 | 0.848406024 | 0.784003812 | 0.736097232 | 0.493501676 |
| 10kb | 0.78391552 | 0.830270283 | 0.921681646 | 1 | 0.933918906 | 0.868093155 | 0.815392267 | 0.584146019 |
| 15kb | 0.737391598 | 0.774522805 | 0.848406024 | 0.933918906 | 1 | 0.941611092 | 0.882449786 | 0.664941136 |
| 20kb | 0.688942755 | 0.721045756 | 0.784003812 | 0.868093155 | 0.941611092 | 1 | 0.944940353 | 0.736770549 |
| 25kb | 0.65370155 | 0.680923471 | 0.736097232 | 0.815392267 | 0.882449786 | 0.944940353 | 1 | 0.789022886 |
| 50kb | 0.445513598 | 0.454410934 | 0.493501676 | 0.584146019 | 0.664941136 | 0.736770549 | 0.789022886 | 1 |

## Code Availability

- Code to compute SItH scores is available upon request: Charles Vaske at.
- Code for all other analysis, including data sets with computed SItH scores, is available at: https://github.com/kjbussey/SItH.

## Data Availability

All data in this study are publicly available for analysis:

- Cancer data: https://dcc.icgc.org/pcawg.
- Normal tissue variant data from the Complete Genomics Indices database in the 1000 Genomes Project (release 20130502): ftp://ftp.1000genomes.ebi.ac.uk/vol1/ftp/release/20130502/supporting/cgi_variant_calls/

## Acknowledgments

We thank Paul Davies, Adam Orr, Charles Lineweaver, Susan Rosenberg, Julia A. Bos and Robert Austin for their insightful discussions into the role of genomic instability and DNA repair in cancer. We thank Sabrina Leung, Terry Christenson and Jessica Pirkle Callan for their technical writing support. We also acknowledge the work of the clinical collaborators, data analysis teams, and funders generating the WGS data in the Pan-Cancer Analysis Working Group of the International Cancer Genome Consortium and the Complete Genomics Indices database in the 1000 Genome Project database.

## Appendix 1 Detailed tuple distributions

We consider a *n-tuple* as a set of *n* contiguous SNVs in genomic space with inter-SNV distances *x* ≤ *D*^⋆^ and *D*^⋆^ = 15 kb. For different values of *n*, we observed numbers of n-tuples in simulated, normal, and cancer data, shown in Figure 1. Singletons (1-tuple) are significantly under-represented for low mutational loads, while tuples of size two or more are typically over-represented with respect to a Poisson point process model.

**Appendix 1 Figure 1.**
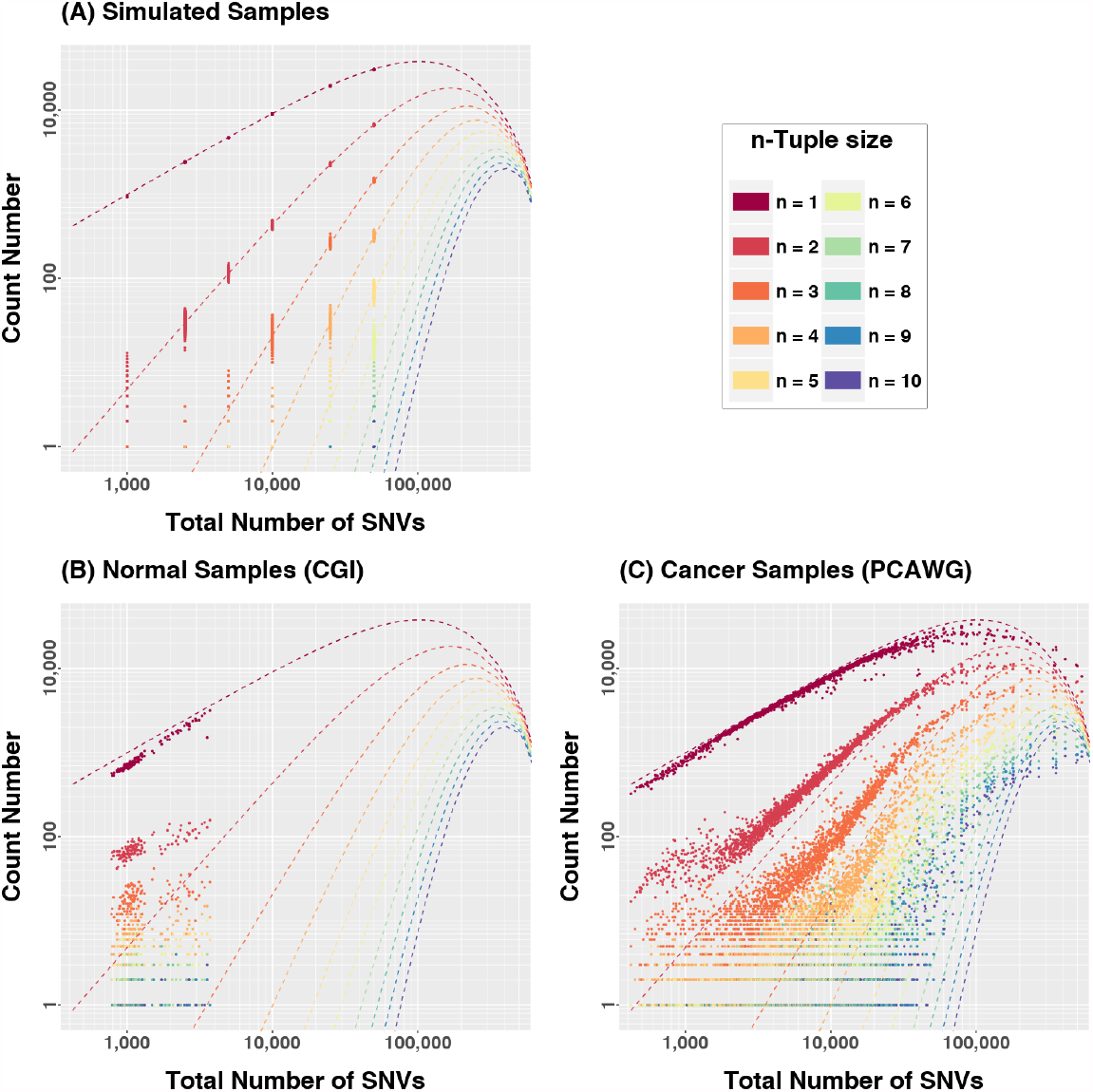
Number of n-tuples in (A) simulated, (B) normal, and (C) cancer samples. Singletons (1-tuples) are less frequent than expected in both normal and cancer cases, while larger n-tuples are more frequent than expected. This relationship is inverted for large mutational loads.

## Appendix 2 Precision in allele fraction estimations

The 95/95 binomial tolerance interval for a true allele fraction of 0.5 at a read depth as high as 60x ranges from 0.25 to 0.75 (Figure 1), meaning that random fluctuations in allele fraction estimations anywhere in that range cannot be ruled out. According to this much larger read depths are necessary to have the precision power to use allele fractions as a method to estimate mutation lineages and discriminate varying degrees of heterogeneity across a tumor.

**Appendix 2 Figure 1.**
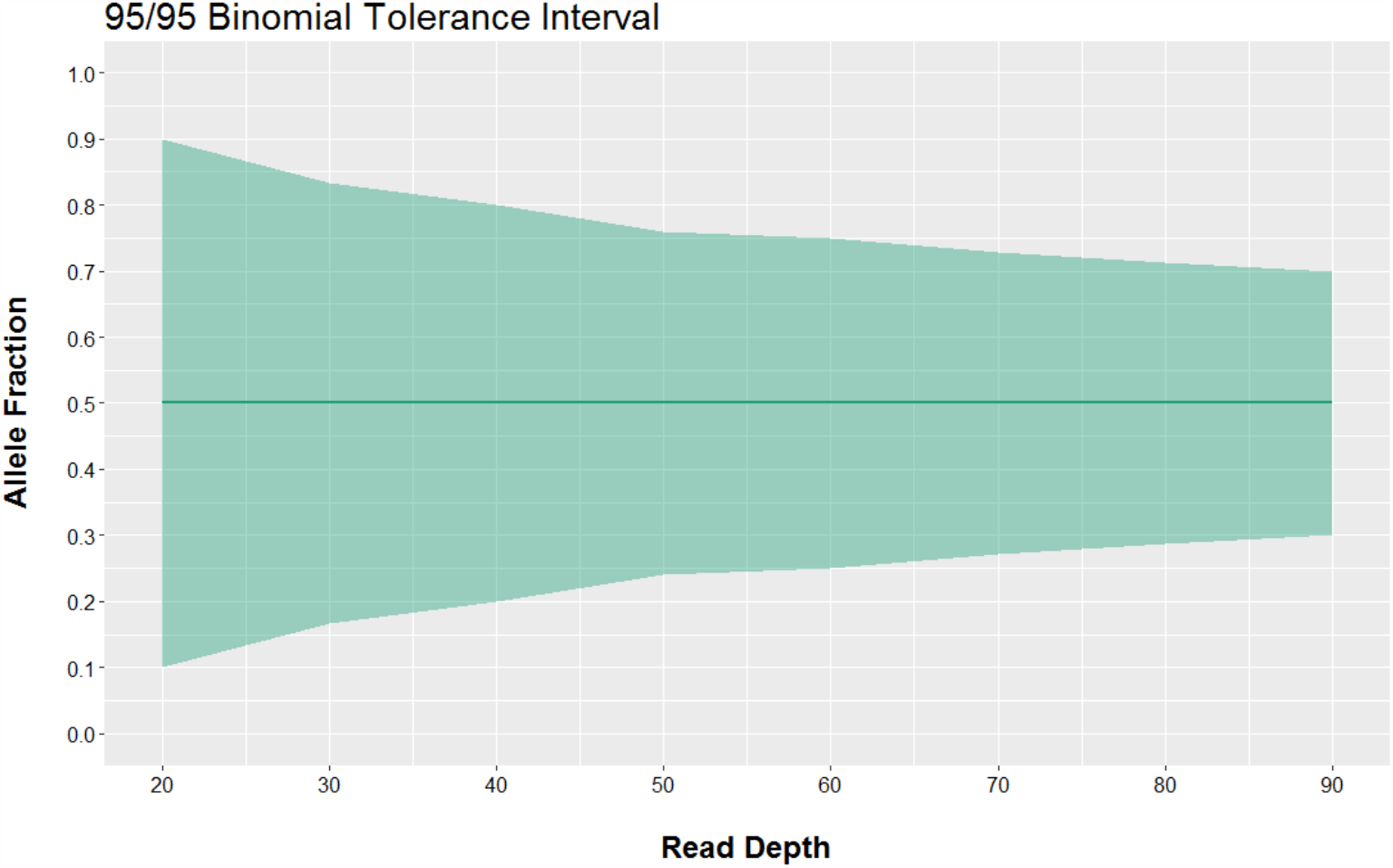
The shaded interval represents the bounds in which we are 95% confident that 95% of the measurements of a true allele fraction of 0.5 will lie as a function of the real read depth.

## Notes

### Competing Interest Statement

The authors have declared no competing interest.

